# Identification of drugs for leukaemia differentiation therapy by network pharmacology

**DOI:** 10.1101/676106

**Authors:** Eleni G Christodoulou, Lin Ming Lee, Kian Leong Lee, Tsz Kan Fung, Eric So, Enrico Petretto, S. Tiong Ong, Owen JL Rackham

## Abstract

Acute leukaemias differ from their normal haematopoietic counterparts in their inability to differentiate. This phenomenon is thought to be the result of aberrant cellular reprogramming involving transcription factors (TFs). Here we leveraged on Mogrify, a network-based algorithm, to identify TFs and their gene regulatory networks that drive differentiation of the acute promyelocytic leukaemia (APL) cell line NB4 in response to ATRA (*all-trans* retinoic acid). We further integrated the detected TF regulatory networks with the Connectivity Map (CMAP) repository and recovered small molecule drugs which induce similar transcriptional changes. Our method outperformed standard approaches, retrieving ATRA as the top hit. Of the other drug hits, dimaprit and mebendazole enhanced ATRA-mediated differentiation in both parental NB4 and ATRA-resistant NB4-MR2 cells. Thus, we provide proof-of-principle of our network-based computational platform for drug discovery and repositioning in leukaemia differentiation therapy, which can be extended to other dysregulated disease states.

## Introduction

The acute leukaemias are a family of haematopoietic disorders which involve blocked differentiation and increased proliferation of immature myeloid or lymphoid blasts in the bone marrow and peripheral blood. The differentiation block and increased self-renewal are consequences of dysregulated gene control, due in turn to the altered activity or expression of transcriptional or epigenetic regulators, as well as signaling pathway components^1^. However, targeting aberrant signaling can be problematic due to mutations in pathway components or bypass mechanisms, which allow for therapy evasion and resistance^2^. An alternative approach is the identification of drugs which can reverse the aberrant activity of transcriptional regulators underlying the differentiation process. Here, we provide this alternative by building a network-pharmacology approach which systematically identifies such drugs.

Transcription factors (TFs) orchestrate the control of gene expression by interacting directly with *cis*-acting DNA sequences, recruiting cofactors as well as epigenetic factors, and in turn initiating or repressing gene expression^3^. Recent studies have shown that leukaemic cell states are reversible and can be manipulated by overexpressing TFs that specify changes in cell fate^4,5^. For instance, a number of “driver” TFs (such CEBPα and PU.1) are well-studied regulators of myeloid lineage specification and differentiation in normal haematopoietic cells^6^. The ectopic overexpression of these TFs in Ph^+^B-cell acute lymphoblastic leukaemia (B-ALL) blasts^4^ or primary chronic myeloid leukaemia (CML) cells in blast crisis^5^ can induce them to undergo myeloid differentiation and lose their leukaemia-initiating potential. In addition, patient-derived AML or CML blasts can be reprogrammed into an iPSC-like state using the Yamanaka TFs (SOX2, KLF4, MYC, and OCT4)^7,8,9^, and subsequently re-differentiated into therapy-sensitive leukaemic cells. However, additional evidence suggests that leukaemias may depend on other TFs to establish the block in differentiation^10,11,12^, and that expression of certain oncogenic TFs may even be subtype-specific^13^. Although such work is enlightening, a structured approach for determining which “driver” TFs provide a regulatory hierarchy over the transcriptional state in specific subtypes is lacking.

We previously developed Mogrify, a computational method which predicts the TFs, along with their networks, required to mediate reprogramming and transdifferentiation between somatic cell types^14^. These driver TFs are ranked based on their influence over the differentially expressed genes within their regulatory network between the two cell states of interest. Here, we envisioned that Mogrify could also be used to identify the best combination of TFs which drive leukaemic differentiation. To demonstrate this, we used Mogrify to identify TFs which facilitate the granulocytic differentiation of acute promyelocytic leukaemia (APL) cell line NB4 upon *all-trans* retinoic acid (ATRA, also known as tretinoin) treatment. We chose this disease model since the process is both thoroughly characterised and robust^15,16^. Mechanistically, ATRA binds to the driver fusion oncoprotein PML-RARα in NB4 cells, changing the repertoire of transcriptional cofactors with which it interacts, thus yielding activation of differentiation genes^17^. Eventually, ATRA induces PML-RARα degradation by proteasomal or caspase-dependent mechanisms^18,19^.

In order to maximise the translational potential of our framework, we implemented a strategy for targeting the selected TF networks using small molecules (including FDA approved drugs). To this aim, we utilised the Connectivity Map (CMAP) repository which contains 6,101 small molecule-induced gene expression profiles which can be compared with externally generated disease signatures^20^. Using these signatures, it is possible to search for a small molecule which can induce specific gene expression changes. CMAP has previously been used for drug repurposing^21,22,23^. By using input signatures specific to leukaemia cells or TFs as the input to CMAP, other studies have identified drugs which eliminate leukaemia cells, induce their differentiation or the loss of self-renewal^24,25,26,27^. Here we extend this approach at the network level, using Mogrify-identified TFs and their networks as the input signature for CMAP, to identify differentiation-inducing drugs. In addition to identifying drugs, our approach provides mechanistic insights into the gene regulatory processes underlying differentiation in APL. Since many drugs in CMAP have been approved by the FDA for other indications, our discovery strategy allows us to avoid the labor, expense, and high failure rates associated with new drug development^28, 29^.

## Results

### ATRA-induces transcriptional changes in NB4 cells

The first key input to our framework are differentially expressed genes in NB4 cells before and after ATRA-induced differentiation. To this end, NB4 cells were treated with 1µM ATRA over a period of 5 days to induce granulocytic differentiation as described previously^15^ (Fig. 1a). Wright’s staining was used to assess differentiation which led to a progressive reduction of the nuclear-cytoplasmic ratio in NB4 cells, whilst the nitroblue tetrazolium (NBT) assay measured a continuous increase in reactive oxygen species (ROS) production. (Fig. 1b, top and bottom panels). These assays showed that ATRA-induced differentiation was essentially complete by day 5.

**Figure 1:**
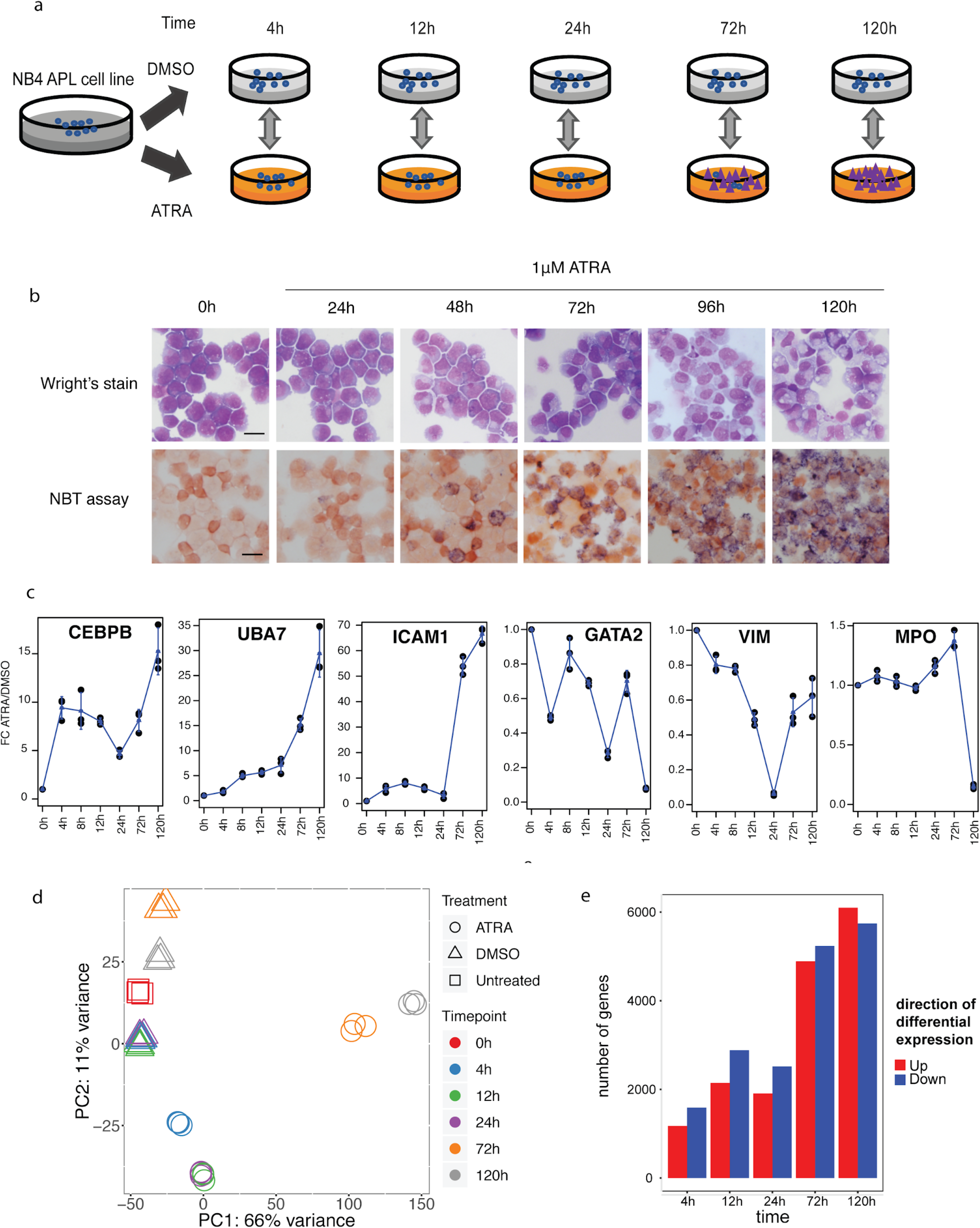
Schema of the experimental design and data analysis of ATRA-induced differentiation of NB4 cells. (a) NB4 cells were treated with 1μM ATRA or DMSO control at five time-points and subjected to RNA-Seq. (b) Wright’s stain and NBT assay of NB4 cells treated with ATRA over 120h. Scale bar, 25μm. (c) qRT-PCR validation of genes known to be up-regulated (top panel) or down-regulated (bottom panel) by ATRA treatment in NB4 cells. Error bars correspond to standard deviations for n = 3 biological replicates. (d) PCA plot of the top 10% most variable genes. (e) Barchart shows the number of differentially expressed coding and non-coding genes between ATRA and DMSO-treated samples. These are coding and non-coding genes and are calculated based on HUGO gene IDs. Suppl. Fig 3 shows the intersections of gene numbers among different time-points. For a full list of DE genes at each time-point, their log_2_ fold changes and their adjusted p-values please see Suppl. Tables 1 and 2.

To characterise these ATRA-induced changes at the transcriptional level, we initially performed qRT-PCR analysis of known differentiation markers at various time-points before and after treatment. Genes were selected from those previously reported to be up- or down-regulated by ATRA in NB4 cells^16,30,31^. We found that these selected genes underwent the expected expression changes, occurring at early (4h: *CEBPB, GATA2*), intermediate (12-24h: *UBA7, VIM*) or late (72h: *ICAM1, MPO*) time-points (Fig. 1c).

**Table 1:** TFs that pass filter 1 and 2, for time-points 4h and 72h.

|  | Up-regulated TFs | Down-regulated TFs |
| --- | --- | --- |
| Time-point 4h | *ID1, HIC1, BCL6, IRF1, CEBPE, ELF4* | *EBF3, IRX3, MYC, MYB, HOXD13, BCL11A, TGIF1, NR2F2, ZEB2, GATA2* |
| Time-point 72h | *EGR3, BATF2, TRIM22, EGR2, IFI16, DDX58, MEF2C, FOXC1, STAT1* | *ZBTB16, SOX18, WT1, CITED1, TBL1X, NFE2, E2F5* |

**Table 2:** Combinations of TFs that pass filter 1, 2 and 3 at time-points 4h and 72h.

|  | Up-regulated TFs | Down-regulated TFs |
| --- | --- | --- |
| Time-point 4h | *ELF4, ID1, IRF1* | *GATA2, MYC, TGIF1* |
| Time-point 72h | *FOXC1, MEF2C, STAT1* | *E2F5, NFE2, WT1* |

To capture the wide-ranging transcriptomic changes induced by ATRA over time, we carried out RNA sequencing (RNA-Seq) on the five most informative time-points: 4h, 12h, 24h, 72h and 120h after treatment, as well as on DMSO controls. We verified that the normalised RNA-Seq counts for the genes of interest were consistent with the qPCR measurements (Supplementary Fig. 1 and 2). Principal component analysis (PCA) of the top 10% most variable genes across time (Fig. 1d) revealed that after only 4 hours of ATRA treatment there was a clear distinction between the ATRA and DMSO-treated cells, with a further drastic change at 72h. Following this, we identified differentially expressed (DE) genes at each time-point (Fig. 1e), which revealed that the most widespread changes occur at 4h and 72h post-treatment, consistent with the PCA.

**Figure 2:**
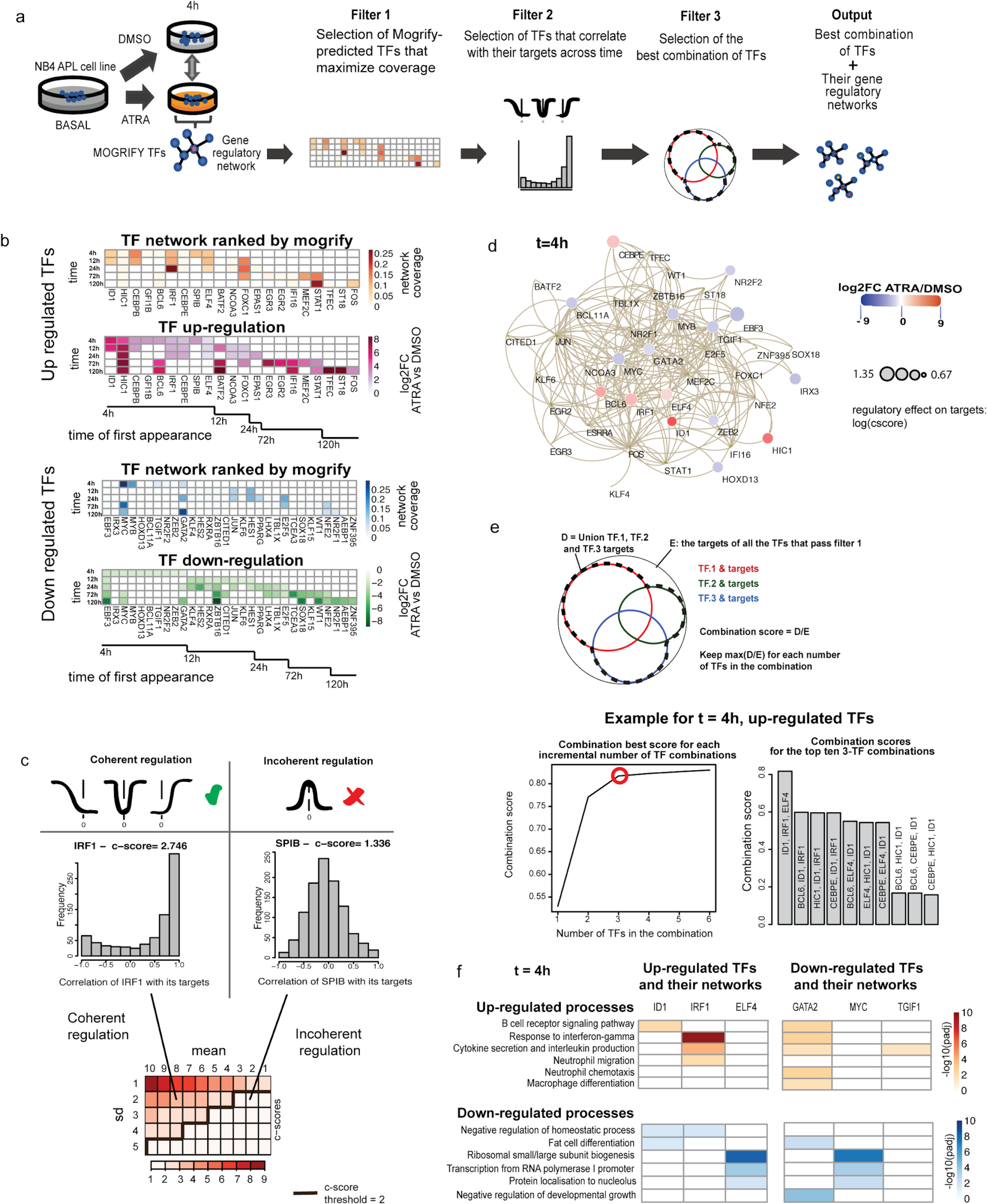
(a) Summary of the pipeline and filters for each time-point: First we detect the driver TFs and their gene regulatory networks. Next, three filters are applied to detect the best combination of TFs whose networks cover the maximum number of DE genes between ATRA and DMSO treated cells. (b) Heatmaps of the network coverage regulated by the Mogrify-selected TFs after applying filter one. The early (4h) and late (72h) wave-like patterns of regulation are evident. (c) Filter 2: Time-wise correlation of expression of selected TFs’ with those of their targets and visualisation of c-score criteria (see Materials and Methods). Good (IRF1) and poor (SPIB) examples of TF-target correlation patterns are shown in the histogram plots. The colored matrix shows the means and the standard deviations of these correlations and the resulting c-score space. Examples of coherent and non-coherent TF-target regulation are shown in the c-score grid. The dark border line on the grid depicts the selected c-score threshold equal to 2. (d) Network of the selected TFs after the application of filter two at 4h. The nodes represent the TFs and their sizes are proportional to the log_2_ c-score, as calculated by filter two. The node color indicates the extent of up- or down-regulation of the TF gene expression in ATRA-treated NB4 cells. The edges represent connections between the node TFs according to the STRING database^33^. (e) Top panel shows a graphical depiction of filter three (see Materials and Methods) with three hypothetical TFs. Bottom left line graph shows the best scores for different numbers of TFs in the combination. Scores represent up-regulated TFs for 4h. The bottom right bar chart shows the scores of the top ten 3-TF combinations for the up-regulated TFs at 4h. (f) Selected GO biological processes and their statistical significance for each TF network of the TF combination, resulting from the application of all three filters at 4h.

To better understand the biological processes involved in ATRA-induced differentiation, we performed Gene Ontology (GO) enrichment analysis of the DE genes (see Materials and Methods). The up-regulated genes detected at early time-points (between 4h and 24h) were enriched for inflammatory and immune system processes (Supplementary Fig. 4, Supplementary Tables 3 and 4), whilst at later time-points we identified a shift towards myeloid differentiation, cell cycle arrest, and apoptosis-specific processes. These ATRA-induced gene expression and biological processes are in agreement with existing literature^16,30,31^, suggesting that our time series captures the important biological features of ATRA-induced differentiation.

**Figure 3:**
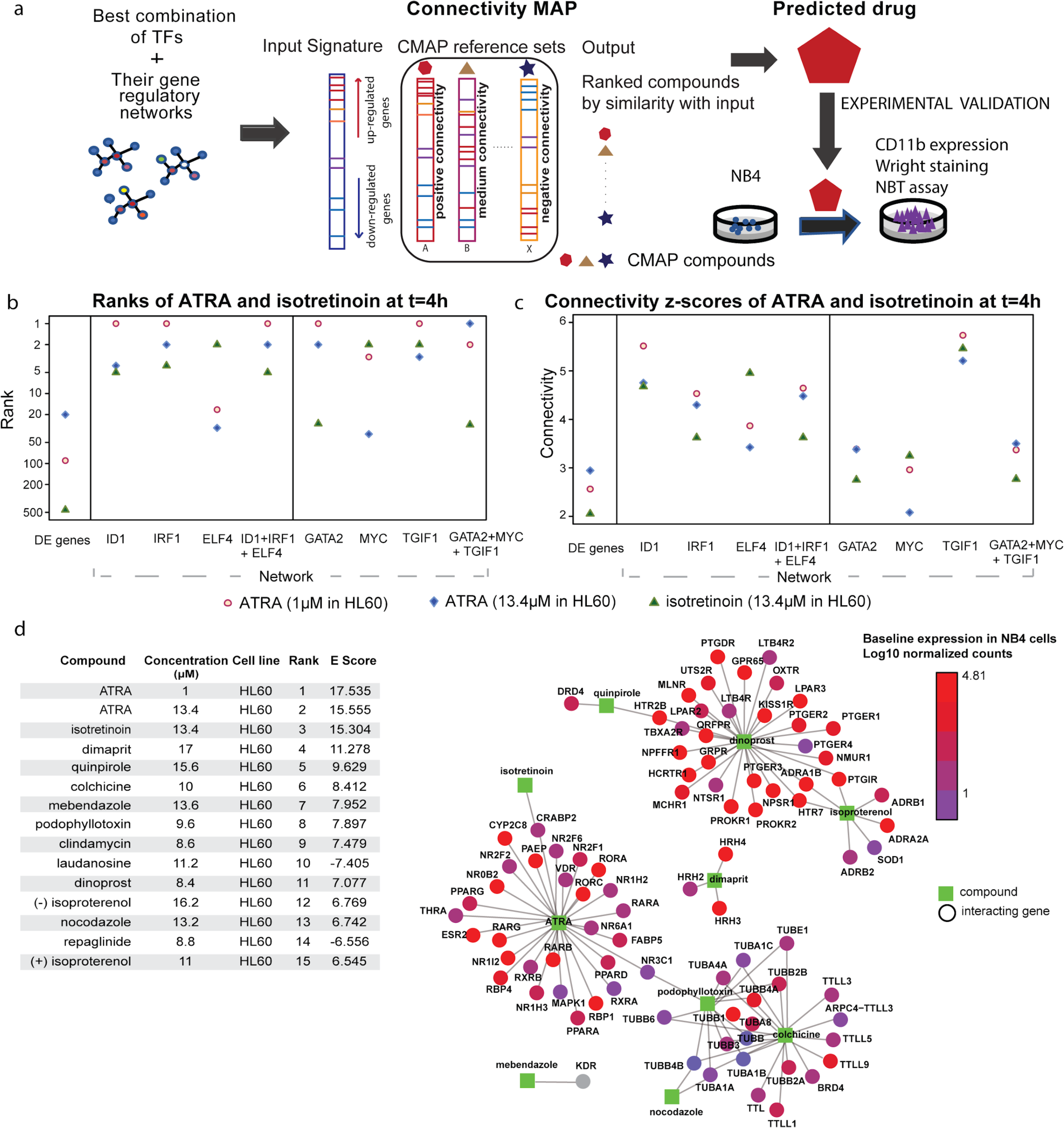
(a) Schematic of the network pharmacology framework. DE genes between ATRA and DMSO-treated NB4 cells were calculated for five time-points using this pipeline. Driver transcription factors (TFs) and their gene regulatory networks were detected using the Mogrify algorithm. This corresponds to the combination of TFs resulting from filter 3 (Fig. 2e). CMAP was queried using these gene regulatory networks. The output of ssCMAP was a ranked list of drugs, from most similar to least similar to ATRA. (b) The ranks of the two ATRA and one isotretinoin cell-line level instances when we ran ssCMAP using different input signatures. (c) The z-scores of the connectivity for the two ATRA and one isotretinoin instances, as calculated by ssCMAP, for different input signatures. (d) Drug-target interaction network of the top 15 drugs (see Supplementary Table 8) as provided by STITCH. Only the targets expressed at 0h (basal time) are depicted. The only exception is mebendazole’s target, KDR, which is not expressed at 0h, therefore it is colored in grey. The table provides detailed drug information ranked by absolute E-Score.

**Figure 4.**
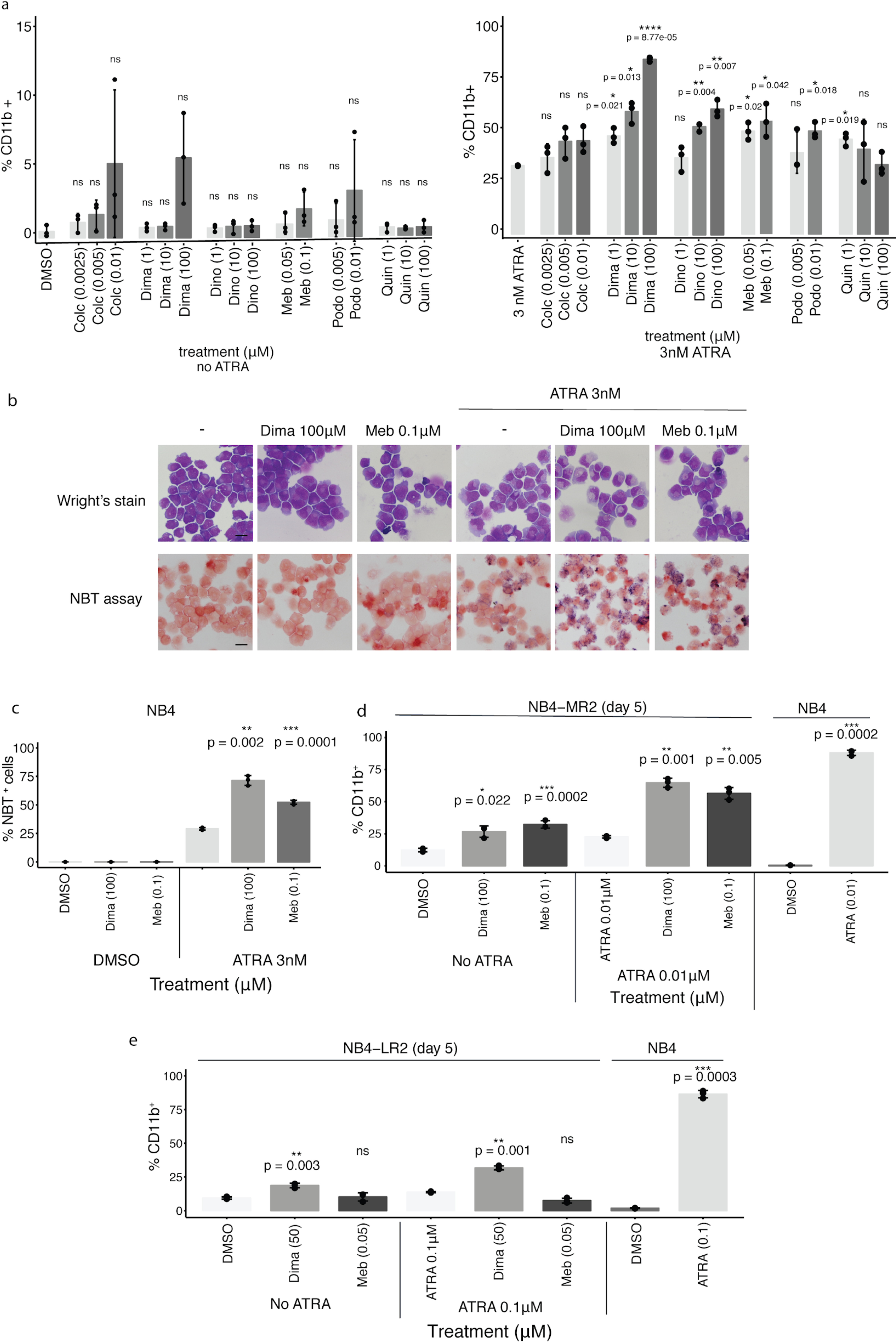
CD11b flow cytometry assay of NB4 cells treated with CMAP positive connectivity drugs colchicine (Colc), dimaprit (Dima), dinoprost (Dino), mebendazole (meb), podophyllotoxin (Podo) or quinpirole (Quin) alone or in combination with a suboptimal dose of ATRA (3nM) for 72h. Error bars represent standard deviation. Enhancement of ATRA-mediated differentiation was observed especially with dimaprit, mebendazole, and dinoprost. ‘ns’ indicates non-significant result (p > 0.05); *p ≤ 0.05, **p ≤ 0.01,***p ≤ 0.001, and ****p ≤ 0.0001 respectively by two-tailed Welch’s t-test (see Materials and Methods). The comparisons are made between each CMAP drug concentration and the respective baseline treatment. (b) Wright’s stain (top) and NBT assay (bottom) of NB4 cells treated for 72h with dimaprit or mebendazole, alone or in combination with ATRA. Scale bar, 20μm. (c) Quantification of NBT assay results in (b) showing that dimaprit and mebendazole enhance ATRA-mediated differentiation. Error bars represent standard deviation with significance represented as in (a). Comparisons are made against treatment with 3nM ATRA alone for each small molecule combination. (d) CD11b flow cytometry analysis of NB4-MR2 cells treated for 120h with dimaprit or mebendazole, alone or in combination with ATRA. Both CMAP drugs overcome ATRA resistance in this cell line. Significance is represented as in (a). The comparisons are made between each CMAP drug concentration and the respective baseline treatment (DMSO or 0.01μM ATRA). (e) Same as (d), but for NB4-LR2 cells. Dimaprit and mebendazole induce minimal differentiation when combined with ATRA. Error bars represent standard deviation with significance represented as in (a). The comparisons are made between each CMAP drug concentration and the respective baseline treatment (DMSO or 0.1μM ATRA).

### Mogrify detects known and novel transcription factors involved in myeloid cell differentiation

We set out to identify TFs and their regulatory gene networks which are induced or repressed by ATRA treatment at each time-point. To this end, we supplemented the Mogrify algorithm^14^ by adopting a multi-step strategy to identify TFs that are (i) predicted to drive ATRA-induced differentiation and (ii) exert a significant transcriptional influence on ATRA-regulated target genes (Fig. 2a).

Initially, we ran the Mogrify algorithm to identify a set of driver TFs, which we then refined in three steps (see Materials and Methods). In the first step, we retained only those TFs which regulate part of the network uniquely (i.e., removing TFs which are redundant to each other) (Fig. 2a, 2b). We identified a wave-like pattern of transcriptional changes induced by the filtered TFs, one at 4h and another at 72h (Fig. 2b), consistent with the observed transcriptional and phenotypic changes (Fig. 1b and 1c), and other reports on ATRA-treated leukaemic cells^16,31^. Therefore, for the rest of our analysis, we focused on these transcriptional waves and investigated perturbations to TFs at the initial and later stages of the differentiation process. We hypothesized that by doing this, we can either simulate the initiating cascade of changes or promote the differentiation once the initial changes have taken place.

The results of filter one show that different TFs may be responsible for different stages of ATRA-induced differentiation. At the early time-points, CEBPB, ID1, GFI1B, GATA2, and MYC may mediate the majority of transcriptional changes, whereas in later time-points STAT1, FOXC1, MEF2C, EGR2, and EGR3 seem more prominent (Fig. 2b). Notably, most of the identified TFs at early and late time-points are known to be involved in the myeloid differentiation cascade, particularly into the neutrophil lineage^32^, which is characteristic of ATRA-induced differentiation.

To further refine the predicted TFs, we next applied our second filter. This filter identifies TFs whose expression is strongly correlated with that of their target genes across time (filter two, see Materials and Methods). Time-wise correlation analysis of the expression profiles of each TF and their targets revealed some TFs whose transcriptional activity is mostly independent of their targets’ expression during the differentiation process (Fig. 2c, Supplementary Fig. 5-8). Therefore, for these TFs, their influence on ATRA-induced transcriptional changes is not coherent across time, and they were removed. Table 1 contains the TFs that passed both filter one and two. Visualisation of the TF-driven gene networks that passed filter one and two revealed that the TFs’ expression and number of targets change substantially across time, revealing critical dynamic responses in the gene expression programs during the differentiation process (Fig 2d, Supplementary Fig. 9, where nodes represent the TFs).

Finally, we investigated the regulatory coverage of each combination of TFs (filter three, see Materials and Methods). We considered “incremental” combinations of TFs, defined as combinations of 2 TFs, 3 TFs, etc. as our ‘combination size’. For each combination, we calculated a combination score (see Materials and Methods) which reflects the number of targets regulated by the TF combination as a fraction of the targets of those TFs that passed filter one at each time-point. We then kept the highest scoring combination of TFs for each combination size (Supplementary Table 7). As shown in Fig. 2e (bottom panel), for the up-regulated TFs at time=4h, the combination score almost reached its maximum when combining up to three TFs. We therefore kept 3 TFs in the TF combinations throughout our analysis, as shown in Table 2 (more details in Materials and Methods).

To better understand the biological processes in which the selected TFs are involved, we performed GO Enrichment analysis of the TF regulatory networks (see Materials and Methods). These networks were enriched for neutrophil migration and chemotaxis, macrophage differentiation and ribosomal small/large subunit biogenesis, processes that are specific to myelopoiesis (see Discussion) (Fig. 2f, Supplementary Fig. 10, Supplementary Tables 5 and 6), suggesting specific and functional specialisation of these TF networks.

### Identification of novel pharmacological inducers of APL differentiation

The aim of this study is to build a computational framework predictive of small molecules, preferably drugs, that can reverse phenotypes associated with leukaemia progression, such as the block in differentiation. These drugs should mimic the key TF-induced transcriptional changes that occur during ATRA treatment of NB4 cells (Fig 3a). To this aim, we accessed the CMAP repository, which contains 6,100 ‘instances’ which capture small molecule-induced gene expression changes in three cancer cell lines, MCF7 (breast cancer), PC3 (prostate cancer) and HL60 (acute myeloid leukaemia/AML)^20^. Each instance is a combination of a drug, a concentration, a cell line and a batch.

There are many methods for extracting drugs that induce a phenotype of interest from CMAP, known as “querying” CMAP. Traditional methods require a user-provided gene expression signature, which they further match with a drug-induced gene expression signature in the CMAP repository. In this study, we queried CMAP using the statistically significant CMAP algorithm (ssCMAP)^34^. Briefly, ssCMAP incorporates statistical significance, weighting of genes and equal contribution of up- and down-regulated genes of the same absolute log_2_ fold change into the CMAP query. Proper querying of CMAP is crucial for accurate drug retrieval; for example, one can average the drug-induced gene expressions across all three cell lines in the repository or focus on cell-line specific results. Considering the different genetic background of the MCF7, PC3 and HL60 cell lines together with the fact that our framework is built on a leukaemia cell line, we decided to implement ssCMAP for cell-line specific results (see Materials and Methods). This is also supported by previous studies on AML using CMAP, which query CMAP at the HL60 cell line level^27,35^. The input to ssCMAP was each TF-driven network or the combined networks of best combinations (see Table 2). We also compared the ssCMAP predictions using the Mogrify predicted TF-networks with those derived from the set of DE genes.

Our positive control ATRA (tretinoin in CMAP) is included in the CMAP repository at two concentrations, 1μM and 13.4μM. Isotretinoin, an ATRA isomer, is also included at 13.4μM. Since our inputs to ssCMAP consist of ATRA-induced gene expression changes, we expected the CMAP ATRA and isotretinoin instances to rank among the top predicted drugs. ssCMAP orders the CMAP instances first by increasing corrected p-value and then by decreasing absolute connectivity. Connectivity is a measure of similarity between the input and the drug signatures, which is positive when the drug has the same effect as the input signature and vice-versa. For time-point 4h, all HL60 ATRA and isotretinoin instances were among the statistically significant results of ssCMAP. In contrast, the ATRA and isotretinoin instances for the MCF7 and PC3 cell lines were not significant.

We compared the ssCMAP ranks of ATRA and isotretinoin predictions when using Mogrify-predicted TF networks or the set of DE genes as the gene expression input to ssCMAP. We found that the Mogrify networks identified either ATRA or isotretinoin among the top predictions (Fig. 3b). In contrast, using the set of DE genes as input, ATRA or isotretinoin were more lowly ranked among the drug predictions, with the highest rank being 20 (rank of 1 is considered the best)(Fig. 3b). In addition, the ATRA and isotretinoin connectivity z-scores, were higher using the TF-driven networks compared to the set of DE genes (Fig. 3c). Therefore, using the Mogrify-predicted TF-networks provided more specific identification of ATRA from the CMAP repository. The results of the drug predictions and the retrieval of ATRA were consistent when other methods were used to interrogate the CMAP repository (Supplementary Text).

We conducted the same analysis using the TF-driven networks (or DE genes) identified at 72h, however, this did not retrieve ATRA or isotretinoin as top rank drugs. This implies that the early 4h TF-driven networks are more informative in capturing the instigating transcriptional changes induced by ATRA, while later time-points may be more reflective of downstream events.

To further proceed with the experimental validation of our predictions, we assigned one ranking score per CMAP instance, which we refer to as the E-score. The E-score combines the individual TF network ranks per CMAP instance (see Materials and Methods). The full E-score ranked lists of predicted drugs at 4h and 72h are provided in Supplementary Tables 8 and 9.

### Identification of CMAP drugs for experimental validation

In order to experimentally validate the predicted drugs identified using ssCMAP, we first filtered the top 15 candidate instances (ranked by E-score) to those with positive connectivity with the TF networks at the 4h time-point (Fig. 3d). Following this, we used the STITCH^36^ database, which contains a curated set of drug and protein-coding-gene interactions, to find the target genes of the top predicted drugs (as described in Materials and Methods). From the targets, we filtered and visualized only those that are expressed at 0h, therefore they are active at baseline allowing the drugs to act on them. Notably, two drugs have no expressed targets at 0h (mebendazole and clindamycin).

Within the resulting drug-target network (Fig. 3d) there are three main classes of proteins interacting with our candidate drugs; Tubulins, Tubulin Tyrosine Ligases (TTLs) and hormonal receptors, such as dopamine receptors (*DRD4*), adrenergic receptors (*ADRA1B, ADR2A, ADRB1, ADRB2*), histamine receptors (*HRH2/3/4*) and nuclear receptors (*RORA, NR2F6, NR6A1, NR1H2*). Tubulins and TTLs have been shown to be effective targets of anti-leukemic chemical agents, such as vinca alkaloids^38,39^. Beyond the identification of ATRA (Fig. 3), this supports the usefulness of TF networks for predicting drugs with known anti-leukemic properties using the CMAP repository.

There is previous evidence that medendazole has anti-leukemic properties in AML^25^ and indirectly interacts with tubulins according to the Therapeutic Target Database (TTD)^37^. Therefore we chose to experimentally test it, although it does not have expressed targets at 0h. To our knowledge, there is no evidence of clindamycin being used to kill or differentiate leukaemia cells, so we did not it experimentally. In the case that there are common targets between two drugs, we kept the one which has additional targets except the shared ones, to avoid testing drugs with similar mechanism of action. For example, we excluded nocodazole because it has two targets, both shared with colchicine which interacts with many more genes. However, if a drug has at least one unique target, we kept it (podophyllotoxin, quinpirole, dimaprit and isoproterenol).

In summary, we selected seven drugs to be experimentally validated based on these criteria: (*i*) they rank high based on E-score (Fig. 3d), (*ii*) they have positive connectivity to the input signature and (*iii*) they have non-redundant targets. Based on these criteria, the seven shortlisted drugs were Prestwick-983 (dimaprit), quinpirole, colchicine, mebendazole, podophyllotoxin, dinoprost (PGF2α) and isoproterenol.

### Experimental validation of CMAP-predicted drugs

To determine if the CMAP-predicted drugs are able to drive or enhance differentiation of NB4 cells, we first sought to identify a working concentration range of our shortlisted CMAP drugs. The results of a cell viability assay (Supplementary Fig. 12) indicated that three drugs (dimaprit, quinpirole, and dinoprost) did not lead to a significant reduction of viable cell numbers (i.e., > 30%) compared with control, even when the drugs were used at high doses (100µM). Isoproterenol was highly toxic beyond 30µM. The remaining three drugs (mebendazole, colchicine, and podophyllotoxin) exhibited a sharp and dose-dependent decrease in viable cell number at sub-micromolar ranges, which was likely due to their primary cytotoxic activity as microtubule inhibitors. In view of this, for the microtubule inhibitors we used drug concentrations at or below IC50 in order to distinguish differentiation effects from cell death.

To determine the effect of each drug on differentiation, we treated NB4 cells for 72h with individual CMAP drugs at doses up to 100µM (for dimaprit, quinpirole or dinoprost), or not exceeding their IC50s (for the microtubule inhibitors). As a readout, we performed a flow cytometry assay of the drugs using the myeloid surface marker CD11b, whose expression correlates with morphological and functional differentiation in NB4 cells^15,40,41^. According to literature^40,41,42,43,44,45,46^, many investigational drugs do not significantly induce differentiation in NB4 cells unless combined with ATRA. We thus hypothesised that PML-RARα could be a critical target, and a core component which inhibits differentiation-inducing pathways or mechanisms. Thus, we also screened NB4 cells with the CMAP drugs alone or in combination with 3nM ATRA, a suboptimal dose which induces minimal differentiation (Fig. 4a; Supplementary Fig. 13a). Isoproterenol was excluded from our screen due to autofluorescence observed even without staining with the CD11b-APC antibody. While none of the remaining six drugs tested significantly induced differentiation on their own, all of them enhanced differentiation induced by 3nM ATRA, with dimaprit and dinoprost showing marked dose-dependent effects. Although quinpirole promoted ATRA-induced differentiation at 1µM, this effect was diminished with higher doses, possibly due to additional non-specific responses to the drug. Altogether, these results suggest that, while our CMAP-identified drugs drive some overlapping transcriptional networks contributing to differentiation, a certain threshold of perturbation with specific targeting of PML-RARα as the centrepiece is required for initiation of the differentiation process.

We further validated the differentiation-enhancing effect of two drugs from our screen using Wright’s staining and the NBT assay. Dimaprit, a cyclic-AMP (cAMP) inducer^47^, was chosen as it was the highest ranking non-retinoic acid (RA) CMAP hit and potently enhanced differentiation. Mebendazole was also selected since it enhanced differentiation at relatively low doses. Furthermore, recent work indicated that mebendazole has anti-leukaemic activity in AML cells^25,48^, although no effects on differentiation were reported. Consistent with the flow cytometry results, neither dimaprit nor mebendazole alone induced differentiation at the morphological or functional levels. However, both drugs caused an increase in the percentage of NBT-positive cells when combined with the suboptimal dose of 3nM ATRA, which induces minimal NBT-positivity on its own (Fig.4b and Fig. 4c, quantification of NBT assay results).

Given these findings, we reasoned that dimaprit and mebendazole would enhance ATRA-mediated differentiation in the NB4-MR2 cell line, which exhibits partial ATRA resistance owing to aberrant corepressor recruitment by PML-RARα^49,50^. Conversely, we predicted that both drugs would be less effective at overcoming ATRA resistance in NB4-LR2 cells, which possess a truncated form of PML-RARα due to a nonsense mutation leading to the elimination of amino acid residues important for ATRA binding^51^. Indeed, we found that both dimaprit and mebendazole induced up-regulation of surface CD11b, particularly when used in combination with ATRA in NB4-MR2 cells (Fig. 4d; Supplementary Fig. 13b). In NB4-LR2 cells, although dimaprit also induced differentiation alone or in combination with ATRA, the effect was less pronounced as compared to that observed in NB4-MR2 cells. Also, mebendazole did not induce differentiation alone or in combination with ATRA in NB4-LR2 (Fig. 4e; Supplementary Fig. 13c). Taken together, these data support that dimaprit and mebendazole rely on at least a partial ATRA response to induce differentiation.

## Discussion

In acute leukaemias, TFs which maintain the differentiation block and increase self-renewal exist in subtype-specific interconnected networks^13^. Perturbations of these networks for therapeutic benefit first require elucidation of components which act as driver transcriptional regulators. In this study, we addressed the use of a data-driven network pharmacology approach for identification of such driver TFs. Using a controlled, time-series RNA-Seq experiment, we predicted putative driver TFs which regulate the ATRA-induced granulocytic differentiation in the APL cell line NB4 using the Mogrify algorithm^14^. We then ranked the relative influence of these TFs before using the highest ranking TFs and their networks (TF-driven networks) as the input for CMAP to identify small molecules which can recapitulate gene-expression activity under ATRA treatment.

We chose to focus on TFs identified at an early stage of differentiation (4h), as they are likely to be initiators of the transcriptional response to ATRA, rather than secondary or tertiary changes^20^. Among the up-regulated TFs, IRF1 and ELF4 regulate expression of genes involved in immune responses, proliferation and/or apoptosis p53^52,53^. In particular, IRF1 is involved in normal granulocytic maturation as well as expression of other crucial regulators of granulocytic differentiation such as CEBPα, ε, and PU.1^54,55^. The other predicted up-regulated TF, ID1, acts as an antagonist of basic helix-loop-helix (bHLH) TFs and has been reported to mediate G0/G1 cell cycle arrest in NB4 cells^56^. Among the down-regulated TFs, GATA2 promotes myeloid progenitor self-renewal^57^ and negatively regulates expression of important TFs which direct granulopoiesis such as CEBPα^58^. Similarly, inhibition of the predicted TF MYC in leukaemia cell lines induced differentiation along various lineages, suggesting that its role in more committed progenitors is to block differentiation^59^. TGIF1 was also predicted as an important down-regulated TF at 4hrs, and is a known repressor of TGFβ signalling which promotes myeloid differentiation^60^. Moreover, TGIF1 inhibits transcription of ATRA target genes^61^. Collectively, Mogrify identified TFs which are involved in directing myeloid development as putative driver regulators of the ATRA-mediated differentiation programme in NB4 cells.

Our results show that, at early responses (t=4h), ATRA is consistently the first or second predicted drug when querying CMAP using the selected TF-driven networks, as opposed to taking the entire repertoire of ATRA DE genes (Fig. 3b-c). The networks of the individual TFs and their 3TF combinations were considered in the final ranking of drugs based on E-Score (Fig. 3d, Supplementary Table 8) (see Materials and Methods). Our hypothesis for including the 3TF combination networks in our drug scoring was that individual TF networks control one part of the differentiation process, but when the networks are combined, they act synergistically and may enhance the entire differentiation process. The ATRA ranks predicted by ssCMAP confirm this hypothesis.

The other drugs (apart from ATRA and isotretinoin), which we predicted using the TF-driven networks in CMAP, induced minimal differentiation on their own, but significantly enhanced differentiation in the presence of ATRA. Unlike other acute leukaemias, APL is considered genetically simple and PML-RARα has been regarded as a crucial driver of disease initiation and of the differentiation block^62^. Thus, we posit that inhibition of PML-RARα with ATRA allows for a level of transcriptional perturbation which permits differentiation responses mediated by our CMAP drugs. In agreement with this idea, previous studies have shown that the ability of various other drugs to induce differentiation in NB4 cells is greatly potentiated by the presence of ATRA. Such drugs include histone deacetylase (HDAC)^41^ inhibitors, plant extract-derived molecules^43^, and cAMP^45,46^.

Among our predicted drugs, dimaprit prominently enhanced ATRA-induced differentiation in a dose-dependent manner and was thus chosen for further validation. Dimaprit is a selective H_2_histamine receptor agonist which was previously shown to increase intracellular cAMP and induce granulocytic differentiation in the HL60 non-APL AML cell line^47^. cAMP induction triggers activation of protein kinase A (PKA), which phosphorylates PML-RARα at S873^63^. This effect was proposed to facilitate proteasome recruitment to PML-RARα and its complete degradation when combined with low-dose RA. This mechanism was used to explain the synergy between cAMP and RA for disease clearance, but not differentiation, in an APL mouse model^63^. In contrast, *in vitro* studies with NB4 cells showed that cAMP enhances differentiation in the presence of retinoid X receptor (RXR) or RARα agonists (such as RA)^45,46,64^. Thus, whether the observed synergy between dimaprit (a cAMP inducer) and low-dose ATRA for differentiation in our NB4 model depends on PML-RARα degradation or transcriptional activation is subject to additional investigation. We cannot exclude the possibility that dimaprit exerts PML-RARα-independent effects, possibly mediated by other genes outside the predicted TFs-driven networks that are concurrently driven by this drug.

Another drug which we selected for further investigation was mebendazole, which is FDA approved for the treatment of parasitic worm infections^65^. We chose this drug due to its efficiency in inducing differentiation (in combination with ATRA) at relatively low doses. Recent studies showed that mebendazole reduces growth and viability of AML cell lines, and induces proteasomal degradation of the oncogenic TF c-MYB through inhibition of its association with the heat shock protein 70 (HSP70) chaperone system^25,48^. However, these studies did not investigate the effects of mebendazole on myeloid differentiation. Here we show that mebendazole enhances differentiation in combination with ATRA in NB4 cells.

Consistent with the notion that the effect of the predicted drugs on differentiation centers on the presence of ATRA, we show that dimaprit and mebendazole can overcome ATRA resistance provided that PML-RARα is also perturbed by ATRA. The NB4-MR2 cell line bears enhanced expression and recruitment of corepressors and chromatin remodelers such as TopoIIβ, NPM1, and BRG1 by PML-RARα, leading to attenuated transcription of RARα target genes^50,66^. These cells still differentiate partially in response to ATRA and are thus susceptible to further maturation induced by dimaprit and mebendazole. In contrast, LR2 cells are virtually non-ATRA responsive. This may be because PML-RARα in these cells contains a nonsense mutation (Gln903*) - mediated C-terminal truncation, leading to deletion of residues important for ATRA binding^51^. Hence, these cells retained their ATRA-resistance even in the presence of dimaprit and mebendazole as ATRA fails to interfere significantly with mutant PML-RARα. Nevertheless, we have not tested the possibility that some combinations of CMAP drugs are able to work on their own without ATRA and are sufficient to override aberrant transcriptional reprogramming caused by PML-RARα. For instance, the top-ranked CMAP drugs at time-point 4h could be used in combination with each other, or with selected top-ranked drugs at time-point 72h to enhance the differentiation process.

The identification of small molecules that can treat acute leukaemias, using specific gene signatures to improve CMAP predictions, is an active field of research. Marstrand *et* al. introduced a ‘conceptual framework’ and identified apoptosis-inducing drugs in APL^35^. In this study, the authors derived APL-specific signatures from public microarray data to query CMAP using the original CMAP method^20^. They predicted a set of drugs which were further experimentally confirmed to induce apoptosis and cessation of tumour cell proliferation. Despite some similarities in the focus on TFs and the use of combinatorial input signatures, our study takes advantage of primary RNA-Seq data on the NB4 cell line and uses mechanistic information to define the optimal and non-redundant set of TF-driven networks to predict drugs using CMAP. This optimization resulted in a higher specificity in identifying ATRA as the top hit (Fig. 3b-c). A few other studies investigating drugs that induce leukaemia differentiation are based on *a priori* knowledge, focusing particularly on the CEBPα transcription factor which directs granulocytic differentiation^67^. In these cases, a CEBPα-induced gene expression signature from the chronic myeloid leukaemia (CML) cell line K562 was used to query CMAP and predict drugs that mimic this gene expression signature^26,27^. These approaches recovered different drugs from the ones we identified, possibly due to (1) the use of microarray data generated in a different cell line for signature derivation, (2) the use of the network of only one TF (CEBPα) and (3) the use of the original GSEA-like method to query CMAP^20^. An important conceptual difference between the aforementioned studies and ours is that we detect the regulatory TFs in a purely data-driven way without using any a priori knowledge. In addition, we further refined the set of non-redundant TF-driven networks by accounting for the co-regulation between TFs and their targets. Finally, the use of curated drug-protein-coding gene interactions (STITCH^36^) allowed us to control for drugs with overlapping mechanisms of action, therefore minimizing redundancy and the number of drugs to be tested experimentally.

In summary, we have developed a network pharmacology framework that prioritises drugs important for leukaemia differentiation therapy. As proof-of-principle, we experimentally demonstrated that the CMAP predicted drugs complement ATRA in driving complete differentiation responses in APL. We showed that the use of a refined input gene signature (here, TF-driven networks) to query CMAP significantly improves drug predictions. In clinical APL, while ATRA treatment induces rapid differentiation of bulk tumour populations, the elimination of self-renewing APL-initiating cells is also required for long-term disease eradication^63,68^. This is typically accomplished through a frontline combination therapy consisting of ATRA and the PML moiety-targeting drug, arsenic trioxide (ATO), resulting in long-term survival rates of above 90%^69,70^. Therefore, a comprehensive differentiation therapy must be coupled to the abrogation of self-renewal in order to be successful. Unlike in APL, where the differentiation block and self-renewal can be attributed to a central oncogenic driver (PML-RARα), it is possible that a multitude of factors, including non-genetic factors, might contribute to these phenotypes in other leukaemia subtypes and diseases. We anticipate that our integrated Mogrify^14^ and CMAP^20^ strategy can prove useful to computationally predict combinations of TFs, their regulated gene networks and drugs which reverse many aberrant disease states, and is not limited to only acute leukaemias and cancers.

## Materials and Methods

### Cell culture and drug treatments

NB4, NB4-LR2 and NB4-MR2 cells (a kind gift from Prof. Eric C.W. So, King’s College London, UK) were cultured in RPMI 1640 (Nacalai Tesque) supplemented with 10% FBS (Biowest), 100U/mL penicillin-streptomycin and 2mM L-glutamine (Gibco). *All-trans* retinoic acid (ATRA, Sigma-Aldrich) was reconstituted in DMSO to 80mM concentration and aliquoted for storage at - 80°C. CMAP drugs were obtained from Sigma-Aldrich with the exception of colchicine (Little Pharmaceutical Suppliers), and reconstituted in DMSO or PBS. Final DMSO concentrations for all experiments did not exceed 0.1% (NB4) or 0.001% (LR2 and MR2). For 5-day experiments, fresh media with drugs was added on the 2nd or 3rd day.

### Wright staining and NBT reduction assay

For Wright staining, 0.5 × 10^5^ NB4 cells were resuspended in 100µL PBS and spun down onto microscope glass slides using a Cytospin™ 4 Cytocentrifuge (Thermo Fisher Scientific) at 500rpm for 5 mins using low acceleration. Slides were dried for 6 min and subsequently incubated at room temperature with Wright stain (Sigma-Aldrich) for 8 mins. Stains were then rinsed off with deionised water and fully dried before mounting cytospots with DPX mountant (Sigma-Aldrich). Brightfield images were taken at 40X magnification using an Olympus IX71 inverted microscope. For the NBT assay^71^, 2 × 10^5^ cells were harvested as before and incubated in 200µL of NBT reaction solution consisting of 1mg/mL NBT (Sigma-Aldrich) and 200ng/mL phorbol myristate acetate (PMA, Sigma-Aldrich) in PBS at 37°C for 30 mins. Cells were then centrifuged onto slides and counterstained with 1% Safranin O (Sigma-Aldrich) in 100% ethanol for 8 mins, dried and mounted as before. A minimum of 100 cells per cytospot were counted for production of blue water-insoluble formazan precipitate indicative of ROS production.

### Quantitative real-time PCR

RNA was extracted using the RNeasy Mini Kit (Qiagen) with on-column DNase digestion. The RNA was further concentrated and purified with the RNeasy Micro kit (Qiagen). RNA integrity was assessed on the Agilent 2100 Bioanalyzer (Duke-NUS Genome Biology Facility, Singapore). Reverse transcription was performed using the Superscript™ VILO™ cDNA Synthesis Kit (Invitrogen). qRT-PCR was performed on the ABI 7900HT Fast Real-Time PCR System (Applied Biosystems) with 2X SYBR Green PCR Master Mix and primers (Table 3). Data were analysed on RQ manager 1.2.1. software and gene expression was normalised against the *TBP* housekeeping control.

**Table 3:**
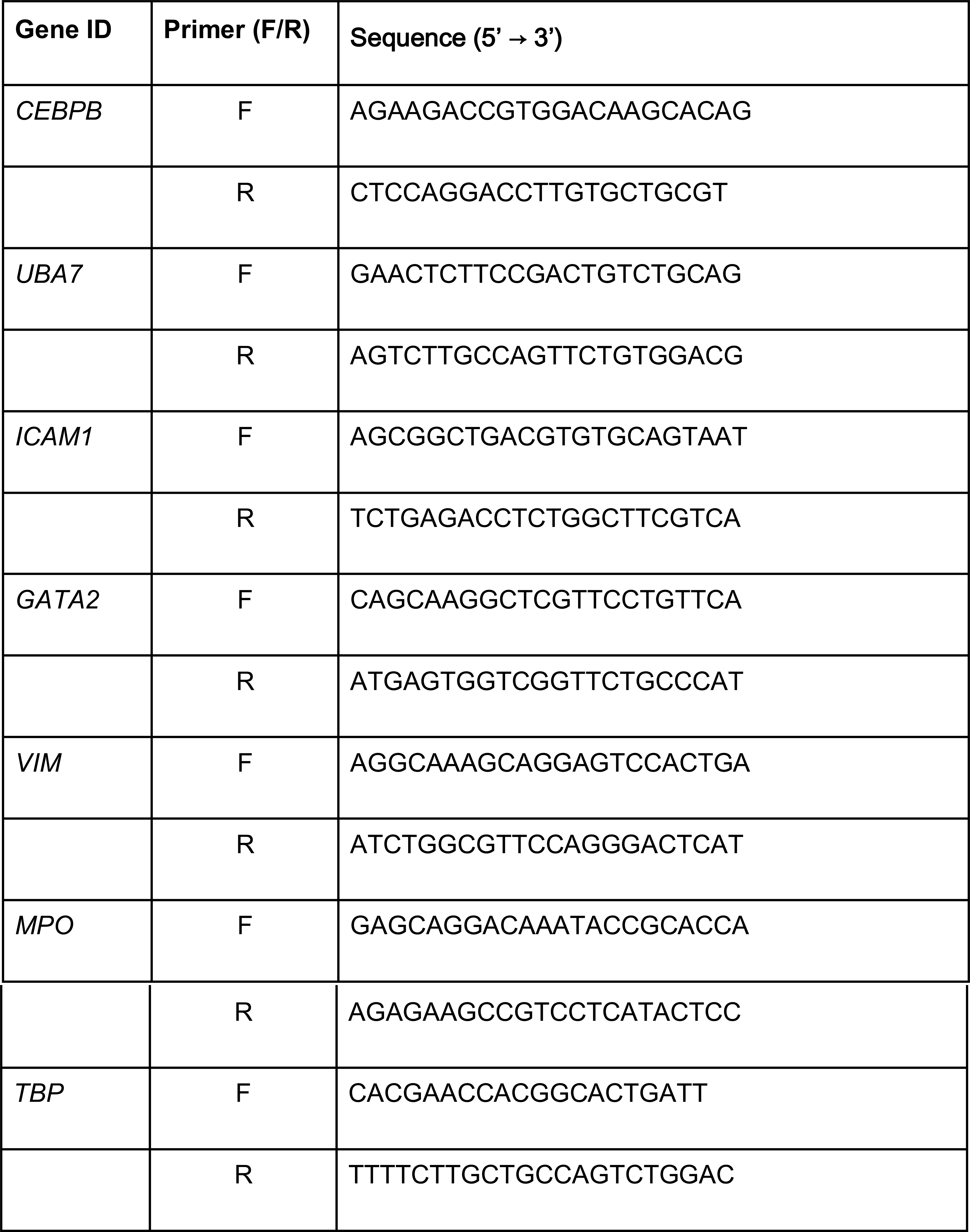
List of primers used in the study.

### Cell viability assay

CellTiter-Glo Luminescent Assay (Promega) was performed according to the manufacturer’s instructions.

### Flow cytometry

Cells were washed once with PBS and stained with CD11b-APC or isotype control (Rat IgG2b-APC, Miltenyi Biotec) antibodies in the presence of human FcR Blocking Reagent (Miltenyi Biotec) at 4°C for 15 mins. Cells were washed again and CD11b surface expression was measured on a BD LSRFortessa™ analyser (BD Biosciences). The data were analysed on the FlowJo v10.5.3. software (TreeStar).

### RNA sequencing

RNA-Seq libraries were prepared from 33 NB4 APL samples using the Illumina Tru-Seq Stranded Total RNA kit protocol, according to the manufacturer’s instructions (Illumina, San Diego, California, USA). Library fragment size was determined using the DNA 1000 Kit on the Agilent Bioanalyzer (Agilent Technologies). Libraries were quantified by qPCR using the KAPA Library Quantification Kit (KAPA Biosystems). Libraries were pooled in equimolar and cluster generation was performed on the Illumina cBOT system (Illumina). Sequencing (150bp pair-end) was performed on the Illumina HiSeq 3000 system at the Duke-NUS Genome Biology Facility, according to manufacturer’s protocol (Illumina). The sequencing depth was at least 37M reads per technical replicate. The data consist of 3 biological replicates, each of which has 2 technical replicates on separate lanes.

### Quality Control, Mapping, Feature counting

The raw RNA-Seq data was subjected to fastqc^72^ and multiqc^73^ protocols, which confirmed that the sequencing was successful and the data were of good quality. Subsequently, three steps were applied:

1. Trimming: Trimmomatic version 0.36 was used to trim the adapters. Trimming was applied on each of the technical replicates at each time-point.
2. Alignment: The technical replicates were merged after trimming. The merged trimmed reads were mapped to the Genome Reference Consortium GRCh38 (hg38) using STAR Aligner v2.5.2b. The Ensembl-90 fasta sequence release was used for indexing. The fasta files for all chromosomes were downloaded from *ftp://ftp.ensembl.org/pub/release-90/fasta/homo_sapiens/dna/* and were concatenated. The paired-end read length found in the STAR aligner log file was used, which was 143bp (STAR was run twice).The reads were further mapped to the human genome using the chimeric STAR option.
3. To quantify the aligned reads of the genes, exons and promoters, the featureCounts module from the Subread suite v1.5.1 was applied.

### Raw Counts Normalisation

The count data were loaded in R-3.4.0 and normalised using the DESeq2 R package v. 1.16.1 ^74^. We used a multifactorial design matrix to model the effect of *treatment* (ATRA or DMSO) and *time*. The time:treatment interaction term was added in the model, to account for the additional changes across treatments due to the effects of time. Only the genes with sum of normalised counts across all samples greater than 10 were kept for further analysis. The PCA plot of Fig. 1b shows the log2-transformed, normalised counts, calculated by the DESeq2 normalisation step.

### Differential Expression

The DESeq2 R package v. 1.16.1^74^ was used for the identification of DE genes between ATRA and DMSO treated samples at each time-point, with DMSO being the reference samples. The design matrix contained the grouping factors time, treatment and time with respect to treatment (∼time+treatment+treatment:time). The resulting log_2_ fold changes were not shrunken. The criteria for differential expression was that the adjusted p-value of the DESeq2 fitting should be less than 0.05. Details on the numbers of DE genes are shown in Fig. 1e and in Supplementary Fig. 3.

### Detection of TFs using Mogrify

The TFs that regulate the DE genes were detected by the Mogrify algorithm^14^. Mogrify predicts TFs that regulate cell conversion by incorporating differential expression and gene network interactions. Mogrify requires as input the set of DE genes at each time-point and their log_2_ fold changes. For each time-point and for each direction of expression (up- or down-regulation), Mogrify calculates the cumulative coverage of the differentially expressed genes by a set of predicted TFs, adding one TF at a time depending on their rank. The following 3-filter process was further applied, to derive the most influential combinations of TFs that control the ATRA-induced transcriptional changes.

Filter 1: From the set of TFs that Mogrify predicted, filter 1 keeps the ones with rank <=10 or target coverage score ≥5%, provided that the rank is below 50. For each TF, its rank is provided by Mogrify and reflects both the number of DE genes it targets and its log2 fold change. The target coverage score was calculated for each TF at each time-point. This was defined as the additional number of genes that the selected TF regulates (compared to the union of all TFs with better ranks) divided by the number of DE genes at the time-point of interest. If a TF was selected at one time-point, it was not re-considered for selection at a later time-point (Fig. 2b)

Filter 2: Correlations of the TFs retained by filter 1 with the expression of their target genes were calculated for all five time-points, using the Pearson correlation metric. The coherence score (c-score) was used to measure the correlation for each TF with the expression of their targets.

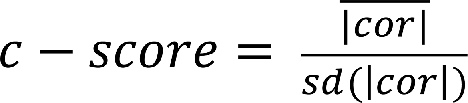

Each TF c-score was calculated considering the TF targets at a specific time-point. The correlation is between the TFs and their target gene expression across all time-points. The TFs with c-score ≥ 2 were passed to the last filtering step. (Fig. 2c, Supplementary Figs. 5-8)

Filter 3: The final filtering step calculates all possible TF combinations, for the respective number of TFs (*x*) that pass filter 2 at each time-point and in each direction separately (up- or down-regulation). For each time-point and for the TF up- and down-regulation separately, let *D* be the union of genes that each TF combination regulates and *E* the number of genes that all TFs that pass filter 1 regulate. For each combination, the combination score *D/E* was calculated and for each *n*-TF combination (for *n* ∈{1, *x*}) the highest score was reported (Fig. 2e, Supplementary Table 7). The best combination scores for 1 to 6 TF combinations of the up-regulated TFs that pass filter 2 at 4h are visualized in Fig. 2e. The combination score was low for 1-TF and 2-TF combinations but increased for 3-TF combinations. Additional TFs did not increase the combination score much further. In order to keep the number of tested TFs to a minimum, the best 3-TF combination was kept for the 4h time-point. For consistency, the best 3-TF combinations for the other time-points of interest were kept as well.

### Gene Sets Functional Enrichments

The R package *gProfiler* was used, querying the GO:Biological Process ontology. For the foreground (query set), separate up- and down-regulated genes were considered. This applies both to DE gene enrichments and to TF network enrichments. For the background set *i)* the full set of expressed genes at the time-point of interest was considered for the enrichment of DE genes (Supplementary Fig. 4) *ii)* the up- and down-regulated DE genes at the time-point of interest were considered for the enrichment of the TF networks (Fig. 2f, Supplementary Fig. 9). The parameters for *gprofiler* function were: max_set_size = 200, min_set_size = 5, hier_filtering = ‘moderate’, max_p_value=0.05. For plotting the GO heatmaps of Fig. 2f and Supplementary Figs. 4 and 9, the GO categories that were expected (based on published literature) or that were highlighted by the REVIGO tool^75^ were selected from Supplementary Tables 3, 4, 5 and 6. Categories that were interconnected, for example cytokine secretion and interleukin signaling, were combined in one category, having as adjusted p-value the most significant adjusted p-value among them.

### Querying CMAP

We queried CMAP using the ssCMAP method in March 2018^34^. The software sscMap.zip.old was downloaded from ftp://ftp.qub.ac.uk/pub/users/sdzhang/PURLsscMap. The software contains the ranks of the genome-wide log_2_ fold changes for the CMAP instances (each instance is a combination of a drug, a cell line, a concentration and a batch). The software at that time had a limitation on the input signature length. We thus ran the ordered version of the algorithm using the top 200 Affymetrix probe IDs of our input signatures, ranked in a software-consistent manner. To convert the HUGO gene IDs of our RNA-Seq experiment to Affy probe IDs, we used the R package *hgu133a*.*db*^*76*^.

CMAP can be queried using ssCMAP at four different levels, which correspond to instances: i) Level 0 (L0): small molecule only, ii) Level 1 (L1) small molecule + concentration, iii) Level 2 (L2) small molecule + concentration + cell line or iv) Level 3 (L3) small molecule + concentration + cell line + batch. Starting from level (iv) and going to level (i), at each step the effects of the small molecule are summarised by the last term. For example, going from level (iv) to level (iii), the effects of the small molecule of the same concentration and of the same cell line are averaged across batches. In our study, we queried at level 2 (L2) small molecule + concentration + cell line. The number of CMAP instances at this level is 3738. ssCMAP calculates Bonferroni-corrected p-values. We executed 10 million permutations in order to have higher distinctive power in the resulting p-values.

### ssCMAP results retrieval and ranking

The first step of processing the ssCMAP outcome was the retrieval of significant results with Bonferroni corrected p-values less than the threshold p-value. By default, only one false positive is allowed, therefore the corrected p-value threshold is:

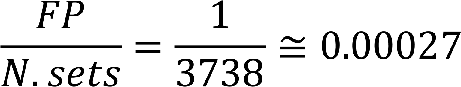

with *FP* being the number of false positives and *N*.*sets* being the number of instances queried at L2. To assign one rank reflective of all the TF network ranks to each drug, we calculated a metric which combines the corrected p-value and the connectivity score of the significant results for each TF network. The connectivity score is a metric introduced in the original ssCMAP publication^34^. It is a number between −1 and 1, indicative of how similar the input and the reference instances are.

When the reference instances constitute a set (of time-points for example) the connectivity score is the average of the individual instance scores. For each drug, the total score was calculated by:

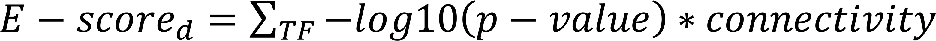

with *d* being the drug of interest. The above formula sums over all the individual and combined TF networks for the TFs in the 3-TF combination. A positive sign indicates positive connectivity and a negative sign indicates negative connectivity. The ssCMAP results were ranked by absolute *E-score* (Supplementary Table 8).

### Detection and network visualisation of drug targets using the STITCH database

The top drugs ranked by E-score were further investigated with the STITCH v5.0 database^36^, before selecting the ones to experimentally validate. STITCH was used to identify the selected drugs’ interacting proteins. The InChI keys of the CMAP drugs were manually retrieved from Sigma Aldrich. The STITCH drugs corresponding to these InChI keys were retrieved. The STITCH entries which contained at least one experimental target and ‘combined score’ > 700 were selected for visualisation by the R *igraph* package^77^ (Fig. 3d). To overlay the expression of the drug targets at 0h, the normalised count values of the respective genes were retrieved from our RNA-Seq experiment.

### CMAP GSEA^78^

Our input sets for GSEA analysis were the ranked small molecules from the CMAP build 2 repository. The respective rank file was downloaded from the CMAP website https://portals.broadinstitute.org/cmap/. This file contains genome-wide gene ranks for each L3 CMAP instance that vary from ∼ 22,000 to 0. Our gene sets (GSEA GMT files) were our 4h TF networks and their best combination resulting from filter 3. For consistency with ssCMAP, the top 200 Affymetrix IDs from the network gene signatures were used to create each gene set.

The Python version of GSEA was implemented with 10,000 permutations. The resulting p-values were adjusted for multiple testing with Bonferroni correction using the R method *p*.*adjust*. The significant small molecule Normalised Enrichment Scores (NES) for the same FDR were then summarised at the cell-line level (L2) for a fair comparison with ssCMAP L2. Finally, small molecules were ranked by this summarised NES following the ssCMAP ranking where they were first ranked by adjusted p-value and then by connectivity.

### Probabilistic CMAP (ProbCmap)^79^

The ProbCmap software was downloaded from http://research.cs.aalto.fi/pml/software/ProbCMap/. ProbCmap calculations are based on the expression of the 930 L1000 landmark genes. Each of our inputs were matched to their intersection with the 930 genes. The more genes in the input, the larger the intersection. Supplementary Table 10 contains the ProbCmap results, but we also examined the effect of the input size on the ProbCmap output. To make the inputs comparable, for each of the TF networks that constituted the input signatures, we considered how many genes it differed from the respective DE genes *D*. We then randomly removed *D* genes from the DE gene signature and ran ProbCmap. We performed 1000 such permutations. We report the frequency of the ATRA (tretinoin) or isotretinoin ranks in these normalised permutations in Supplementary Fig. 11.

### Ccmap

The Ccmap^80^ R package was originally developed for querying CMAP with public disease expression data obtained from the NCBI Gene Expression Omnibus^81^. In our context, it was modified to work with an input of a custom set of genes. The full networks of TFs and the full list of DE genes with HGNC gene IDs constituted the input to this algorithm.

### Statistics

Two-tailed Welch’s t-test with confidence interval 95% was applied on the three replicate measurements to calculate the significance of the flow cytometry results (Fig. 4).

## Supporting information

Supplementary Table 10 - referred to in the Supplementary Text

## Data availability

The raw and processed data were submitted to NCBI GEO. The accession number will be available upon publication.

## Code availability

The source code of our pipeline will be available upon publication.

## Acknowledgments

The authors would like to acknowledge Ms. Sonia P Chothani for providing the RNA-Seq data analysis pipeline, the ssCMAP developer, Dr. Shu-Dong Zhang, for helpful discussion on the algorithm’s internal calculations and the Duke-NUS Genome Biology Facility (DGBF) for sample RNA sequencing. This work was funded by the National Medical Research Council (NMRC) of Singapore (MOH-000059/MOH-CSASI18may-0002) and NMRC/CIRG/1429/2015 to S.T.O.). O.J.L.R is supported by NMRC YIRG (NMRC/OFYIRG/0022/2016).

## Author contributions

Wet lab experiments were carried out by L.L.M. Data were analysed and interpreted by E.G.C., O.J.L.R. and E.P. E.G.C. and L.L.M prepared the manuscript and generated the figures with input from co-authors. K.L.L, O.J.L.R., E.P. and S.T.O. designed the study. E.S and T.K.F. provided the NB4 cell line.

