## Supplementary Text for "Identification of drugs for leukaemia differentiation therapy by network pharmacology"

**Application of different CMAP querying methods**

There is a plethora of methods developed for querying CMAP with a custom gene expression signature^1^. The original CMAP querying method is based on a GSEA-like algorithm. This approach handles the reference signature genes as a set and does not consider their ranks. In addition, it handles the up- and down-regulated genes separately in both the input and the output signatures and ignores the magnitude of differential expression. We chose ssCMAP to query CMAP because it incorporates ranking and weighting of genes, without handling the up and down regulated genes separately. In addition, it is one of the few methods that can provide results at the cell line level and not averaged at the compound level. The majority of existing methods for querying CMAP compare gene expression profiles using GSEA-like or pattern matching algorithms. To compare with the ssCMAP findings, we repeated our predictions with two such methods, GSEA^2^ and ccmap^3^, and with one probabilistic method, probabilistic connectivity mapping (ProbCmap)^4^.

These methods are designed to capture the compound rankings at the summarized drug level (L0), therefore they ignore all cell-line, concentration and time-related specific effects. An intervention to original GSEA was needed to extract significant compounds at the cell-line level (L2) and be able to compare directly with ssCMAP (see Methods for details). For ccmap and ProbCmap such an intervention was not possible because the algorithms are not open-source.

All the applied methods retrieved ATRA (tretinoin) among the top hits, with very few exceptions when the TF networks were used as input signatures. Supplementary table 10 contains the top 50 results of the applied methods, highlighting the tretinoin or isotretinoin instances. For each predicted drug by one of the methods, we provide the drug’s ranking by the rest methods. The rest of the drugs that we predicted were often retrieved by these additional methods, but in different rank order. The rest of the results varied among methods. This discrepancy is expected because the methods and the length of the gene expression input differ. The compared methods were not conclusive on whether the DE genes constitute a less accurate input for CMAP than the network, as we concluded using ssCMAP; they rather detected tretinoin among the top hits both for the networks and the DE genes inputs. This could be due to the summarization of results at the compound level and to inherent method design.

**Results of different CMAP querying methods**

Querying CMAP with the modified for L2 GSEA (cell-line level) and for the TFs of t=4h detected tretinoin as first hit for IRF1 and for the combination ID1, ELF4 and IRF1, 3rd for DEgenes, 18th for ID1 (mebendazole was the first) but it was not significant for ELF4. GATA2 and TGIF1 also picked tretinoin among their top hits.

ccmap at L0 identified tretinoin or isotretinoin among the top 5 hits for the networks and for the DE genes input. Full gene signatures were used as input to the method.

ProbCmap internally compares the input gene expressions to the 930 landmark gene expressions of L1000 across 718 compounds. The original number of CMAP compounds is 1309. The extent of intersection of the input signature with the landmark genes is crucial for the outcome of ProbCmap. 186 of the DE genes intersect with the 930, whereas the individual network coverage ranges from 17 for ID1 to 85 for GATA2 and MYC. ProbCmap also identified tretinoin as first hit for DE genes but also for GATA2, IRF1 and combination of 3 TFs networks. Despite lower coverage in the case of individual networks, tretinoin was among the top 6 hits for all the input networks, except for ID1, where it ranked 18th. Tretinoin’s high ranks using individual networks indicate the high accuracy of the network signature. The lower rank of tretinoin in ID1 result is explained by the low intersection between ID1 network and the L1000 landmark genes. In an effort to make the inputs comparable, we employed a permutation approach (see Methods) and report the number of times that tretinoin ranked in the top 6 positions (supplementary figure 10).

ssCMAP was also applied at the summarized drug level (L0). At this level, tretinoin was not the first hit neither for the network or the DE genes input, indicating the importance of the CMAP instances summarization level.

It is worth noticing that not all drugs are tested on HL60 cell line; only 1229 out of the 6100 instances of CMAP build 2 correspond to HL60 cell line. Therefore, when querying at L0 there is a bias in favor of drugs that are universally tested when summarizing across cell lines.
